# CD46 targeted ^212^Pb alpha particle radioimmunotherapy for prostate cancer treatment

**DOI:** 10.1101/2022.10.14.512321

**Authors:** Jun Li, Tao Huang, Jun Hua, Qiong Wang, Yang Su, Ping Chen, Scott Bidlingmaier, Allan Li, Zhongqiu Xie, Anil Bidkar, Sui Shen, Weibin Shi, Youngho Seo, Robert R. Flavell, Daniel Gioeli, Robert Dreicer, Hui Li, Bin Liu, Jiang He

## Abstract

We recently identified CD46 as a novel prostate cancer cell surface antigen that shows lineage independent expression in both adenocarcinoma and small cell neuroendocrine subtypes of metastatic castration resistant prostate cancer (mCRPC), discovered an internalizing human monoclonal antibody YS5 that binds to a tumor selective CD46 epitope, and developed a microtubule inhibitor-based antibody drug conjugate that is in a multi-center phase I trial for mCRPC (NCT03575819). Here we report the development of a novel CD46-targeted alpha therapy based on YS5. We conjugated ^212^Pb, an *in vivo* generator of alpha-emitting ^212^Bi and ^212^Po, to YS5 through the chelator TCMC to create the radioimmunoconjugate, ^212^Pb-TCMC-YS5. We characterized ^212^Pb-TCMC-YS5 *in vitro* and established a safe dose *in vivo*. We next studied therapeutic efficacy of a single dose of ^212^Pb-TCMC-YS5 using three prostate cancer small animal models: a subcutaneous mCRPC cell line-derived xenograft (CDX) model (subcu-CDX), an orthotopically grafted mCRPC CDX model (ortho-CDX), and a prostate cancer patient-derived xenograft model (PDX). In all three models, a single dose of 20 μCi ^212^Pb-TCMC-YS5 was well tolerated and caused potent and sustained inhibition of established tumors, with significant increases of survival in treated animals. A lower dose (10 μCi ^212^Pb-TCMC-YS5) was also studied on the PDX model, which also showed a significant effect on tumor growth inhibition and prolongation of animal survival. These results demonstrate that ^212^Pb-TCMC-YS5 has an excellent therapeutic window in preclinical models including PDXs, opening a direct path for clinical translation of this novel CD46-targeted alpha radioimmunotherapy for mCRPC treatment.

**Significance:** This study reports a novel CD46 targeted ^212^Pb alpha particle radioimmunotherapy, ^212^Pb-TCMC-YS5, that is well tolerated and shows potent anti-tumor activity (tumor growth inhibition and increase of animal survival) *in vivo* in three prostate cancer small animal models, i.e., a subcutaneous and an intraprostate orthotopic mCRPC cell line-derived xenograft models, and a prostate cancer patient-derived xenograft model. Given that YS5 is a clinical stage human antibody, this YS5-based ^212^Pb alpha particle therapy has potential of translation to the clinic for treatment of mCRPC patients.

## Introduction

Targeted radiotherapy with α-particle emitting radionuclides is a promising approach to eliminate microscopic clusters of malignant cells due to the short path length (50–80 μm), high linear energy transfer (LET; 100 keV/μm), and high relative biological effectiveness of α-particles (1). α-particles are known to induce cell death or proliferation arrest regardless of the cell’s oxygen levels or sensitivity to chemotherapy or low LET (0.1–1 keV/μm) radiotherapy treatment (external beam or β-particle radioimmunotherapy (RIT)). ^212^Pb has a half-life (t_1/2_) of 10.6 h that acts as an *in vivo* generator for the high LET α-emitting daughter isotopes ^212^Bi with t_1/2_=60.6 min and ^212^Po with t_1/2_=0.3 microseconds. RIT with ^212^Pb-TCMC-trastuzumab has been clinically tested in ovarian cancer, which showed an antitumor activity and was well tolerated (2,3). Several preclinical studies and a recent clinical trial with ^212^Pb also showed promising therapeutic efficacy in different cancers (4–8).

Current efforts on targeted radio-ligand delivery in prostate cancer are centered on prostate-specific membrane antigen (PSMA) for the adenocarcinoma subtype using anti-PSMA antibodies (e.g., ^177^Lu-J591 and ^225^Ac-J591), or radio-labeled small molecule PSMA-617 (e.g., ^177^Lu-PSMA-617 (9,10), ^225^Ac-PSMA-617 and ^212^Pb-PSMA-617 (11,12)). In 2022, the FDA approved Pluvicto™ (^177^Lu-PSMA-617 or lutetium Lu 177 vipivotide tetraxetan) for the treatment of adult patients with PSMA-positive mCRPC who have been treated with androgen receptor pathway inhibition and taxane-based chemotherapy (13). Despite these exciting development of PSMA targeted radio-ligand therapies, there are several limitations: (1) Radio-labeled PSMA-617 causes persistent xerostomia in patients, impairing quality of life (14); (2) PSMA is heterogeneously expressed, especially in treatment emergent small cell neuroendocrine prostate cancer (15). Not surprisingly, the success of current PSMA-targeted RITs is predicated on patient stratification using PET that detects high levels of PSMA expression (10).

We have previously developed a non-gene expression-based approach to identify tumor cell surface epitopes including those formed by conformational change and post-translational modifications (16,17). We selected billion-member human antibody phage display libraries on patient samples with the aid of laser capture microdissection and identified tumor-binding antibodies following counter-selection on normal tissues (17). Following further characterization including tissue specificity (17) and in vivo tumor targeting by imaging methods (18), we used these tumor selective antibodies to pull down target antigens and establish their molecular identity by mass spectrometry analysis. We discovered a new lineage independent cell surface antigen CD46 that is homogeneously expressed in both adenocarcinoma and small cell neuroendocrine subtypes (15). We further developed a novel human monoclonal antibody YS5 that binds to a tumor selective epitope on CD46 and generated an antibody-drug conjugate (ADC) by conjugating auristatin derivatives to YS5 (15). The ADC is in a multi-center phase I trial (NCT03575819) and a phase I/II combination trial (NCT05011188) for mCRPC. Our recent work also demonstrated that the ^89^Zr radiolabeled YS5 (^89^Zr-DFO-YS5) can detect prostate cancer *in vivo* in both PSMA positive and PSMA negative mCRPC cell line xenograft (CDX) models and patient derived xenograft (PDX) models of both the adenocarcinoma and neuroendocrine subtypes (19), and a first-in-human study of this novel PET agent is ongoing in mCRPC patients (NCT05245006).

In this study, we report the synthesis and evaluation of tolerability and therapeutic efficacy of the ^212^Pb-radiolabeled human antibody YS5 (^212^Pb-TCMC-YS5) in multiple prostate cancer small animal models, including subcutaneous CDX, intraprostate orthotopic CDX, and prostate cancer PDX models.

## MATERIALS AND METHODS

### Reagents

Hydrochloric acid (Trace Metal Grade) was purchased from Fisher Chemical. Pb-resin^™^ (100– 150 μm) was purchased from Eichrom Technologies LLC (Lisle, IL USA). The fully human CD46-targeted monoclonal antibody, YS5, was produced and purified as described previously (15). The radiolabeling chelator, p-SCN-Bn-DFO (1-(4-isothiocyanatophenyl)-3-[6,17-dihydroxy-7,10,18,21-tetraoxo-27-(N-acetylhydroxylamino)- 6,11,17, 22- tetraazaheptaeicosine] thiourea) (catalog No. B-705), and p-SCN-Bn-TCMC (S-2-(4-Isothiocyanatobenzyl)-1,4,7,10-tetraaza-1,4,7,10-tetra(2-carbamoylmethyl)cyclododecane) (catalog No. B-1005) were purchased from Macrocyclics, Inc. ^89^Zr oxalate was purchased from the Cyclotron Laboratory at University of Wisconsin, Madison (Madison, WI). The ^224^Ra/^212^Pb generators used for these studies were purchased through the Isotopes Program, US Department of Energy, Oak Ridge National Laboratory (ORNL; Oak Ridge, TN USA). All other chemical reagents and materials were the highest grade available from Sigma-Aldrich and used as received without further purification (unless otherwise described).

### ^89^Zr radiolabeling of YS5

Conjugation of YS5 with the chelator DFO and subsequent ^89^Zr labeling of DFO-YS5 were performed as described (19).

### ^212^Pb radiolabeling of YS5

YS5 was first conjugated with the bi-functional chelator 2-(4-isothiocyanotobenzyl)-1,4,7,10-tetraaza-1,4,7,10-tetra-(2-carbamoylmethyl)-cyclododecane (TCMC; Macrocyclics, Plano, TX) at the molar ratio of TCMC/YS5 at 10:1. The average TCMC chelator/mAb ratios were determined by a spectrophotometric assay (20). ^212^Pb radiolabeling of YS5-TCMC and purification of the ^212^Pb-TCMC-YS5 were performed as previously described (21) with modifications. Briefly, ^212^Pb was eluted with 2 M HCl from the ^224^Ra-^212^Pb generator and pre-concentrated by passing a column loaded with the Pb-resin. Following the pre-concentration step, about 2 mCi to 5 mCi ^212^Pb^2+^ was eluted from the Pb-resin column into a reaction vessel with pH = 6 NaOAc buffer as the radiolabeling reaction. The reaction vessel was pre-loaded with 200 μg to 500 μg YS5-TCMC in 500 μL pH = 5.4 NaOAc buffer and 50 μL human serum albumin (10 mg/ml) and the reaction vessel was heated at 37 °C for 1 h. The reaction mixture was run on a PD-10 column for purification with 0.1 M phosphate buffer with human serum albumin (10 mg/ml). Radiolabeling efficiency (the percentage of incorporated radionuclide from radiolabeling reaction) and radiochemical purity of ^212^Pb-YS5 was analyzed using radio-HPLC. HPLC analysis was performed using a Zenix-C-SEC-300 (Sepax Technologies, Inc.) with 0.1 M phosphate buffer (pH 7.2) at the eluting rate of 1 ml/min.

### *In vitro* cell binding affinity

The apparent K_D_ value of ^212^Pb-TCMC-YS5 binding to the CD46-expressing mCRPC cell line PC3 was determined by a saturation binding assay with concentrations at 5, 10, 25, 50, and 100 nM as previously described (18).

### Animal subjects and husbandry

CDX studies were performed using 5–7 week old male athymic nude mice (Nu/Nu Nude mice, Charles River, Wilmington, MA). PDX studies were performed using 6-8 weeks old male NSG mice (NOD/SCID/IL-2Rγ^−/-^) (The Jackson Laboratory, Bar Harbor, ME). Animal studies were approved by the University of Virginia Institutional Animal Care and Use Committee and performed in compliance with guidelines from the Public Health Service Policy and Animal Welfare Act of the United States.

### *In vivo* ^89^Zr-DFO-YS5 PET imaging and biodistribution studies

Approximately 3–5 weeks after tumor implantation, animals with PC3 xenografts reaching 100 mm^3^ in volume were anesthetized by isoflurane inhalation. For ^89^Zr-DFO-YS5 PET imaging, 3.70–5.55 MBq (100–150 μCi, 30 μg/mouse) of ^89^Zr-DFO-YS5 in saline was administered through the tail vein, and animals were imaged at 24, 72, and 120 h post injection, with a 20-minute acquisition time by using microPET/CT Albira Trimodal PET/SPECT/CT Scanner (Bruker Corporation). PET imaging data were acquired in list mode and reconstructed using Albira Software Suite provided by the manufacturer. The resulting image data were then normalized to the administered activity to parameterize images in terms of %ID/g. Imaging data were viewed and processed using an open source Amide software. CT images were acquired following PET, and the CT data were used for attenuation correction for PET reconstruction, and anatomic reference.

### Dose study of ^212^Pb-TCMC-YS5

To determine the safe dose for therapy study, a dose escalation study was performed in nude mice without tumor to determine toxicity. Groups of nude mice (n = 5-7/group) were given doses of ^212^Pb-TCMC-YS5 at 0, 5, 10, 20, and 50 μCi per mouse. Mice were monitored for health status, signs of stress and body weight twice weekly. Mice were sacrificed when loss of body weight was more than 20%, or if the mice showed visible signs of distress, including lack of appetite, excessive lethargy, or formation of sores and rashes.

### Blood analysis for hematology and clinical chemistry

To evaluate the potential toxicity of ^212^Pb-TCMC-YS5, hematological analysis and clinical chemistry were performed as reported (5). CD-1 mice (Charles River Laboratories, Wilmington, MA) were administered with 20 μCi of ^212^Pb-TCMC-YS5 as study group. The control group received no treatment. Blood was collected via the orbital sinus into EDTA coated tubes and analyzed immediately using the Heska Element HT5 hematology analyzer (Barrie, Ontario, Canada). For blood chemistry, plasma was separated from blood by centrifugation (12,000 rpm, 10 min, at 4 °C) and was either studied immediately or stored at −80 °C for later analysis. Biomarkers for organ function such as albumin, blood urea nitrogen (BUN), creatinine, alanine aminotransferase (ALT) and aspartate aminotransferase (AST) were measured using enzyme-linked immunosorbent assay (ELISA) kits (ELISA kits for albumin, blood urea nitrogen, creatinine, alanine aminotransferase were purchased from MyBioSource, San Diego, CA; and ELISA kit for aspartate aminotransferase were purchased from Abcam, Boston, MA).

### *In vivo* ^212^Pb-TCMC-YS5 therapy studies in a subcutaneous PC3 xenograft model

For the therapeutic study, mice bearing PC3 tumors about 100 mm^3^ in volume were given a single i.v. dose of ^212^Pb-TCMC-YS5 at 20 μCi (740 kBq) (n = 5) or the unlabeled YS5 antibody (n = 7) or no treatment (n = 4). Tumor dimensions were measured and recorded 3 times weekly with a digital caliper, and tumor volumes were calculated using the following modified ellipsoid equation: volume = 0.5 x length x width x width (19). Tumor growth was calculated as mean percent change in volume relative to the initial volume at the time of dosing. Animal survival data was analyzed by Kaplan-Meier analysis (GraphPad). Body weight was measured twice weekly, and the mice were monitored for signs of distress. Mice were sacrificed if any one of the situations occurred: tumors reached 2.0 cm in diameter, loss of body weight more than 20%, or if the mice showed visible signs of distress, including lack of appetite, excessive lethargy, or formation of sores and rashes. The same health/humane endpoint applies to all our animal studies (subcu- and ortho-CDXs and PDX).

### *In vivo* ^212^Pb-TCMC-YS5 therapy studies in an intraprostate orthotopic PC3-Luc model

PC3-luc cells were derived from PC3 cells transfected with the firefly luciferase expressing plasmid from Addgene (Ubc.Luc.IRES.Puro, RRID:Addgene 33307) (22). PC3-luc cells (3 × 10^5^) were injected into the murine prostate in 30 μL of PBS, as previously described (23–25). Tumor growth was monitored weekly by *in vivo* bioluminescent imaging (BLI) using a Lago X (Spectral Instruments Imaging, Tucson, AZ, USA). Images were acquired 10 min after intraperitoneal injection of 3 mg d-Luciferin Potassium Salt (Gold Biotechnology, Inc., St Louis MO, USA) in 200 μL 0.1 M phosphate buffered saline. Signals were quantified using Aura (Spectral Instruments Imaging) with the manual ROI tool to determine the amount of photons emitted for a given time.

For the therapeutic study, 30 days after PC3-luc cells were inoculated and tumor engraftment were confirmed by *in vivo* BLI, the mice were given a single i.v. dose of ^212^Pb-TCMC-YS5 at 20 μCi (740 kBq) (n = 4) or the unlabeled YS5 antibody (n = 5). Tumor growth was monitored by *in vivo* BLI. Animal survival data was analyzed by Kaplan-Meier analysis.

### *In vivo* ^212^Pb-TCMC-YS5 therapy study in a prostate cancer PDX model

PDX mice with subcutaneously grafted tumor (Cat#: TM00298 PR1996) were purchased from The Jackson Laboratory (Bar Harbor, ME USA). When tumors size reaches about 100 mm^3^, mice were given a single i.v. dose of ^212^Pb-TCMC-YS5 at 20 μCi (740 kBq) (n = 6), 10 μCi (370 kBq) (n = 8) or the unlabeled YS5 antibody (n = 4), each along with non-binding human IgG (0.5 mg per mouse) for blocking Fc receptors. Tumor dimensions were measured twice weekly with a digital caliper. Tumor growth was calculated as mean percent change in volume relative to the initial volume at the time of dosing. Animal survival data was analyzed by Kaplan-Meier analysis. Body weight and signs of distress were assessed twice weekly.

### Statistical analyses

Two-tailed Student’s t-test and one-way analysis of variance (ANOVA) were used for analysis of two and multiple (more than two) groups, respectively. P < 0.05 was considered significant. Survival was analyzed by Kaplan-Meier analysis. The significance of median survival (treatment group vs. saline control) was determined by a Log-rank (Mantel-Cox) test.

## Results

### Radiochemistry and *in vitro* cell studies

The chemistry for ^212^Pb radiolabeling of the YS5 antibody was performed as shown in **Fig. 1A**. The ^212^Pb radiolabeling process was adopted from the Pb-resin pre-concentration and column extraction method described previously, with modifications (21). This process simplifies the handling of the isotope, and shortens the whole process to only 1.5 – 2 h with a labeling yield around 70%. The radiochemistry purity of the final ^212^Pb-TCMC-YS5 was > 98% by radio-HPLC (**Fig. 1B**). On average each YS5 antibody bears about 2 TCMC molecules as determined by a spectrophotometric assay (20). The apparent binding affinity (apparent K_D_) ^212^Pb-TCMC-YS5 to PC3 cells was determined by an *in vitro* cell binding assay and found to be 21.7 ± 7.3 nM (**Fig. 1C**).

**Fig. 1.**
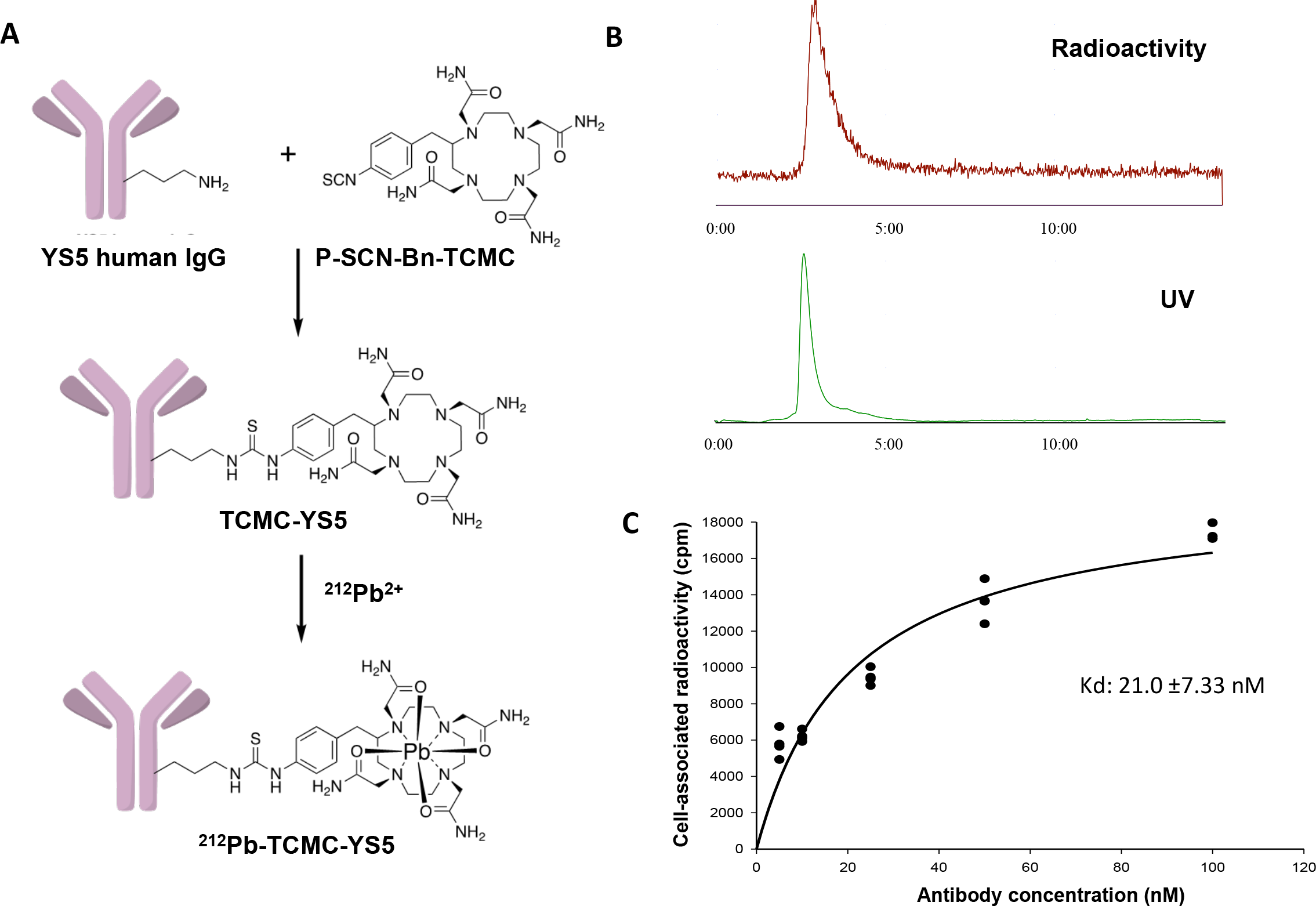
Synthesis of ^212^Pb-TCMC-YS5 and *in vitro* cell binding study. (A) Reaction scheme for ^212^Pb-TCMC-YS5 synthesis. (B) Size-exclusion HPLC analysis of ^212^Pb-TCMC-YS5. (C) *In vitro* cell binding assay of ^212^Pb-TCMC-YS5 on PC3 cells. Triplicates. Apparent binding affinity (K_D_) was calculated by curve fitting.

### Dose and toxicity study

**W**e performed a dose escalation study to identify the safe dose range using non-tumor bearing nude mice. The study included 4 dose groups (n = 5 per group) of 5 μCi, 10 μCi, 20 μCi, 50 μCi and one control group receiving “cold” YS5 (radioactivity defined as 0 μCi, n = 7). As shown in **Fig. 2A**, at the highest dose of 50 μCi, there was 80% death and profound loss of body weight within 10 days. Other groups receiving 5 μCi, 10 μCi, or 20 μCi exhibit dose-dependent body weight loss within the first two weeks followed by a rebound and stabilization throughout the investigation period and have a 100% survival rate (**Fig. 2B**). The control group showed gain of body weight in a slow but steady manner and no death. From this data, the tolerated dose range is 5 – 20 μCi, and 20 μCi was chosen as the dose for therapeutic study in tumor-bearing mice.

**Fig. 2.**
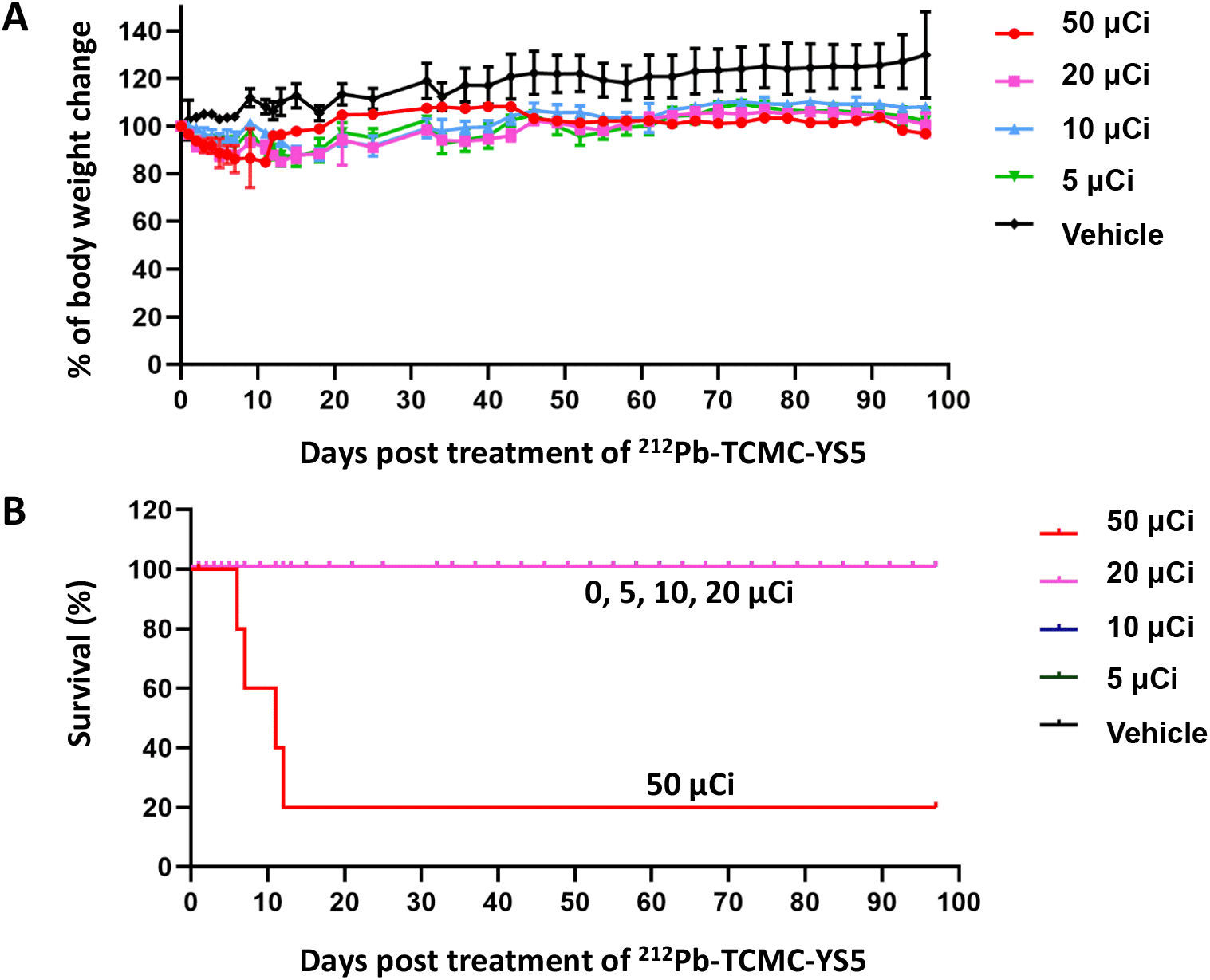
^212^Pb-TCMC-YS5 dose selection study in non-tumor bearing nude mice. (A) Body weight changes post ^212^Pb-TCMC-YS5 treatment. (B) Survival of mice post treatment.

### Hematological and clinical chemistry study

We performed additional studies, blood cell count and blood chemistry, to evaluate both acute (day-14 post treatment) and long-term (day-90 post treatment) toxicity in mice receiving 20 μCi of ^212^Pb-TCMC-YS5. As shown in **Fig. 3A** and **Supplemental Table S1**, there is no significant difference between treatment vs. control group on either day-14 and Day-90 for any of the cell types studied, except for neutrophil that showed a significant increase on day-14 for the treatment group, suggesting an acute but not long-term neutrophil reaction to treatment. The blood chemistry analysis from albumin, creatinine, BUN, AST, ALT were in the normal range and exhibit no significant difference between the treatment vs. the control group (**Fig. 3B** and **Supplemental Table S2**). Those data suggest that 20 μCi of ^212^Pb-TCMC-YS5 is well tolerated and an appropriate dose to use for therapy study.

**Fig. 3.**
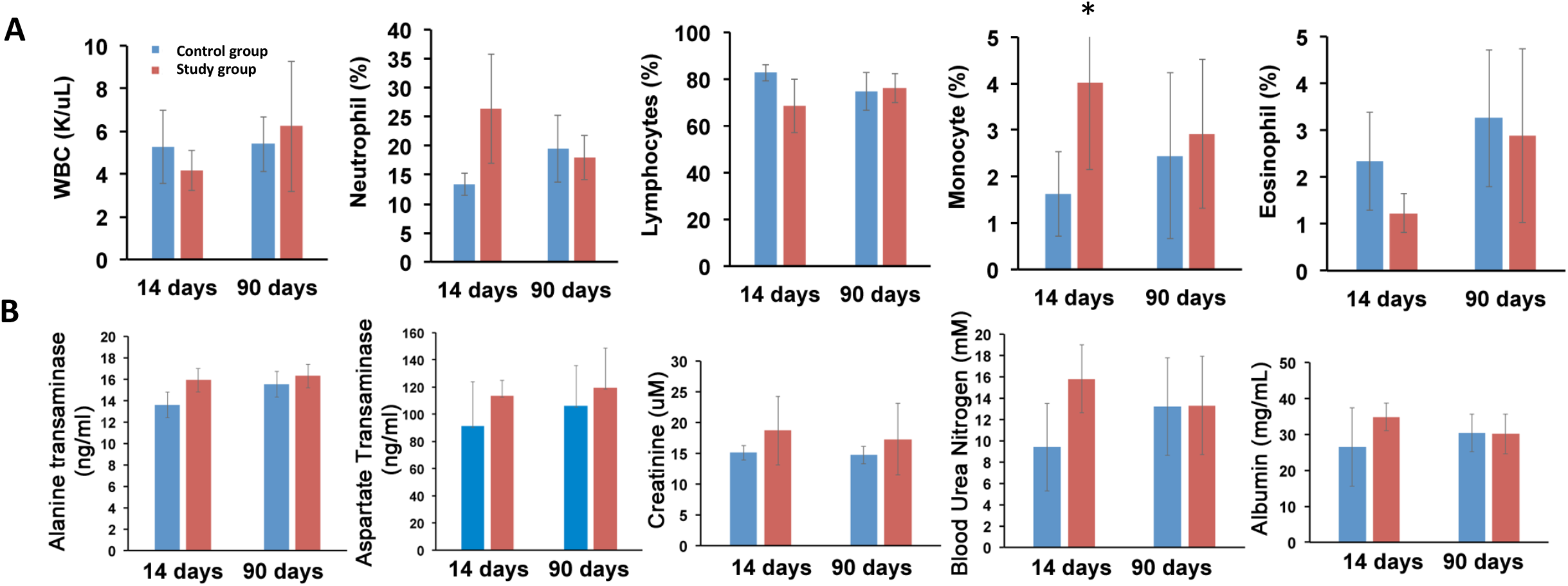
Acute and long-term toxicity assessment by blood analysis. CD-1 normal mice received 20 μCi ^212^Pb-TCMC-YS5 (study group) or no treatment (control group). (A) Hematology analysis of mice 14 days and 90 days post-treatment. (B) Blood clinical chemistry analysis of mice 14 days and 90 days post-treatment.

### Therapy study in a subcutaneous mCRPC CDX model

We first studied anti-tumor efficacy of ^89^Zr-DFO-YS5 in the PC3 subcu-CDX model. We showed, by PET imaging (**Fig. 4A**) and biodistribution (**Fig. 4B**), that ^89^Zr-DFO-YS5 targeted the PC3 tumor *in vivo*, with similar tumor uptake and tumor/non-target ratios compared with other prostate cancer CDX models in our previous study (19). We performed a single dose efficacy study of ^212^Pb-TCMC-YS5 using the PC3 subcu-CDX. A significant tumor inhibition in all treated mice was observed (100%, a complete response): tumor reduction began within the first week with a volume decrease up to 70%, and the inhibitory effect lasted over 40 days post administration of ^212^Pb-TCMC-YS5 (**Fig. 5A**, with individual mouse data shown in **Fig. 5B**). Tumor growth resumed in 3/5 mice after 40 days post treatment (**Supplemental Fig. S1**). Only a transient body weight reduction was observed over the duration of the experiment (**Fig. 5C**), consistent with the result of the dose escalation study in non-tumor bearing mice. All mice recovered to their initial weights within 2 weeks post treatment. 80% mice in the treatment group survived 55 days post treatment (**Fig. 5D**). In contrast, the control groups (cold YS5 antibody and untreated) showed rapid tumor growth (**Fig. 5A**) and 100% death within 37 days (**Fig. 5D**). There were no significant differences between the two controls.

**Fig. 4.**
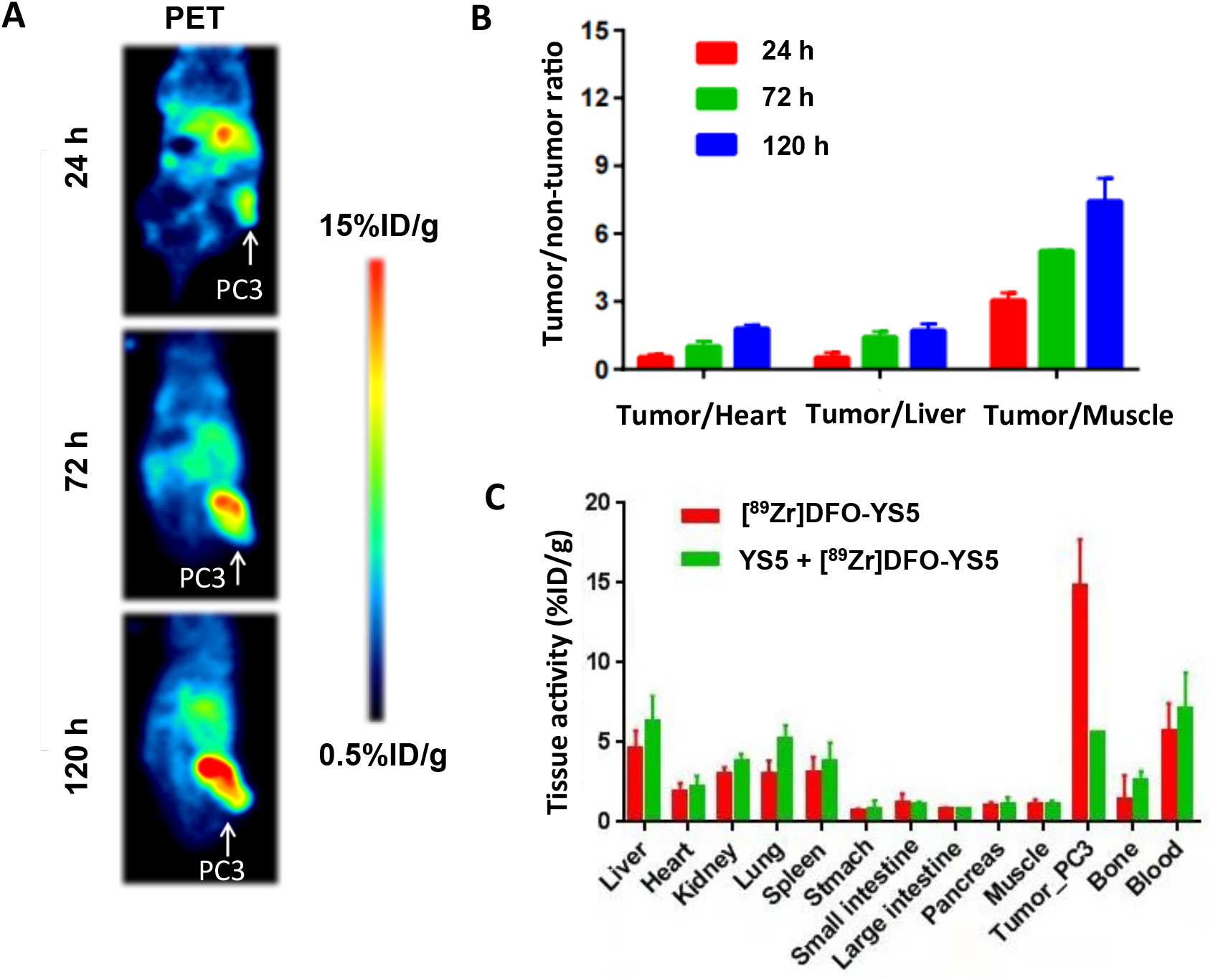
PET imaging and biodistribution of ^89^Zr-DFO-YS5 in PC3 subcu-CDX model. (A) PET/CT images of PC3 tumors at 24, 72 and 120 h post-administration of ^89^Zr-DFO-YS5. (B) Tumor/non-tumor ratio derived from PET images at 24, 72 and 120 h post-administration of ^89^Zr-DFO-YS5. (C) Biodistribution of ^89^Zr-DFO-YS5 at 120 h post-administration.

**Fig. 5.**
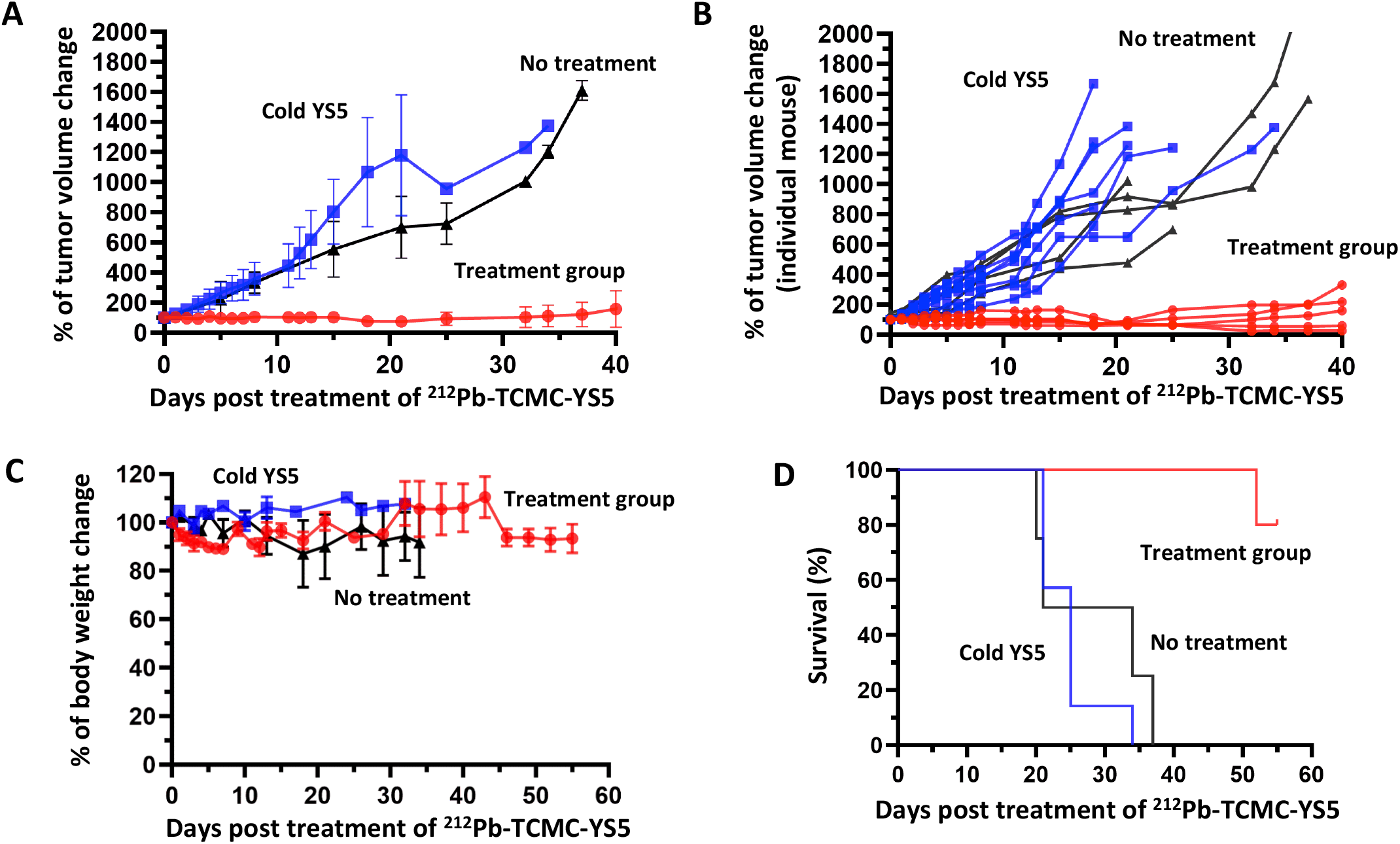
Therapeutic effect of a single dose of ^212^Pb-TCMC-YS5 in the PC3 subcu-CDX model. Red: study group receiving ^212^Pb-TCMC-YS5. Blue: control group receiving cold YS5 antibody only. Black: no treatment. (A) Tumor growth within 40 days post treatment of ^212^Pb-TCMC-YS5 (averaged tumor volumes normalized against tumor volume at day 0). (B) Tumor growth of individual mouse xenografts within 40 days post treatment of ^212^Pb-TCMC-YS5 (normalized against tumor volume at day 0). (C) Body weight post treatment of ^212^Pb-TCMC-YS5 (averaged body weights normalized against body weight at day 0). (D) Survival of mice post treatment of ^212^Pb-TCMC-YS5.

### Therapy study in an intraprostate orthotopic mCRPC CDX model

We further studied the therapeutic effect of a single dose ^212^Pb-TCMC-YS5 in an intraprostate orthotopic mCRPC CDX (**Fig. 6A**). The PC3 cell line expressing the firefly luciferase reporter gene was used to create the ortho-CDX model. As shown in **Fig. 6B**, tumor growth, as measured by BLI (representative images shown in **Supplemental Fig. S2**), was significantly inhibited up to 70% and the duration of inhibition was over 60 days for the study group treated with 20 μCi ^212^Pb-TCMC-YS5, while tumors in the control group continuously grew. Body weight remained steady for the treatment group for the duration of the study (141 days post treatment) (**Fig. 6C**). Survival data is shown in **Fig. 6D**. The mice in the study group survived up to 141 days post treatment (end of experiment) with no death within the first 60 days post treatment. In contrast, no mice survived for the control group 36 days post treatment. The median survival was 100.5 days for the study group, which is significantly longer than the control group that is only 23 days (p = 0.005).

**Fig. 6.**
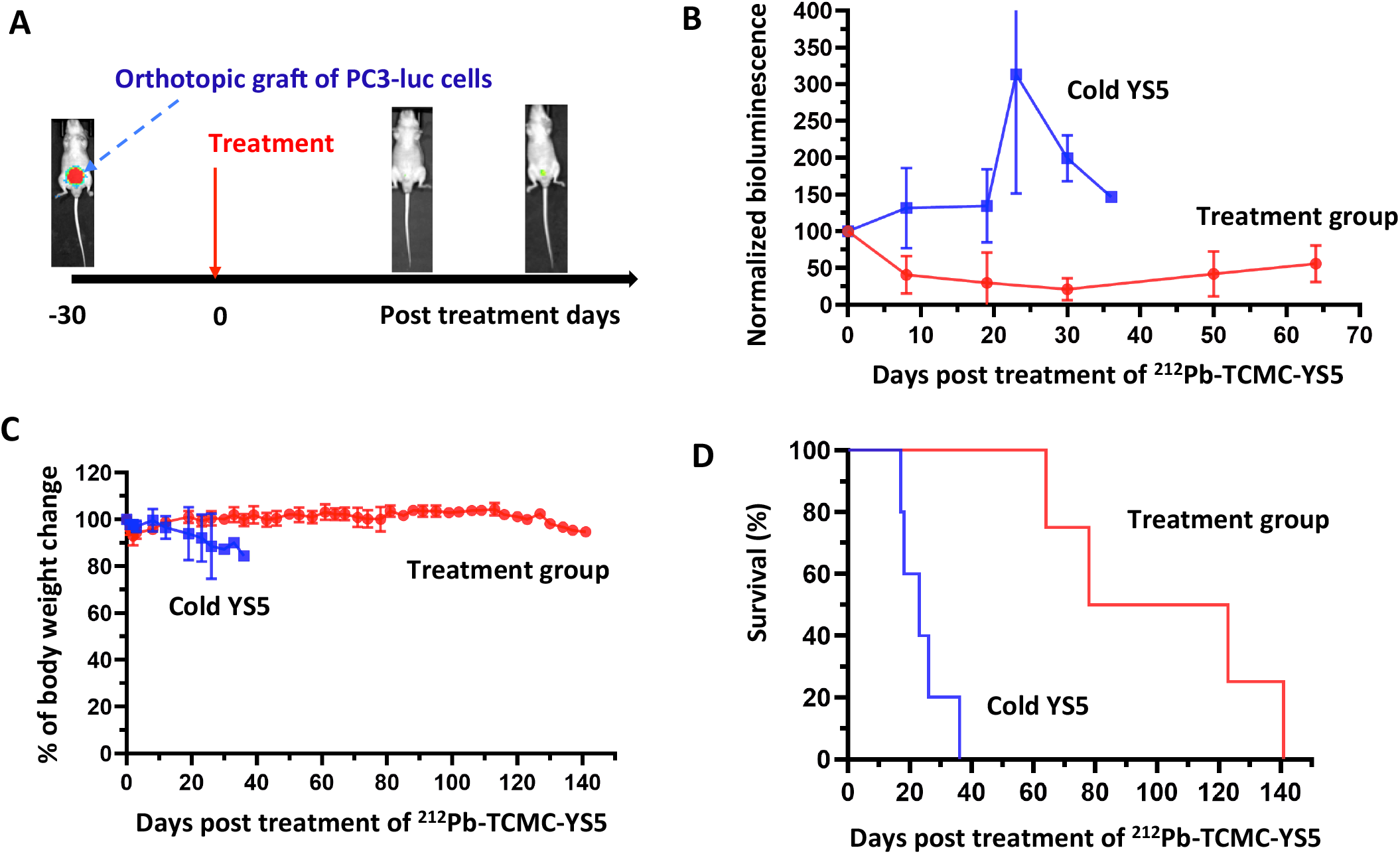
Therapeutic effect of a single dose of ^212^Pb-TCMC-YS5 in the intraprostate PC3-Luc ortho-CDX model. (A) Illustration of study design. (B) Tumor growth measured by BLI. Red: treatment group. Blue: Cold YS5 control.

### Therapy study in a subcutaneous prostate cancer PDX model

To broaden clinical applicability, we performed therapy study using a prostate cancer PDX model. We first used PET imaging with ^89^Zr-DFO-YS5 to confirm tumor targeting by YS5 *in vivo*. The result showed a clear evidence of tumor targeting by YS5 (**Fig. 7A**) with tumor uptake above 20%ID/g from PET images (**Fig. 7B**). Uptake in other organs and tissues was minimal, consistent with our previous study using different PDX models (19). We next studied the therapeutic efficacy of ^212^Pb-TCMC-YS5. The PDX tumor was implanted subcutaneously, allowing ready tumor size measurement by caliper. We studied two doses, 20 μCi, and the lower dose of 10 μCi. As shown in **Fig. 8A** (normalized tumor volume) and **Supplemental Fig. S3** (actual tumor volume of individual PDX), both doses inhibited tumor growth, with the higher dose of 20 μCi showing a longer duration of inhibition. Body weight remained steady for the two treatment groups (10 and 20 μCi) for the duration of the study (**Fig. 8B**). The median survival was at 34 days, 56 days and 99 days respectively for the control, 10 μCi, and 20 μCi groups (**Fig. 8C**), showing a significant improvement over the control for both treatment groups (p < 0.001).

**Fig. 7.**
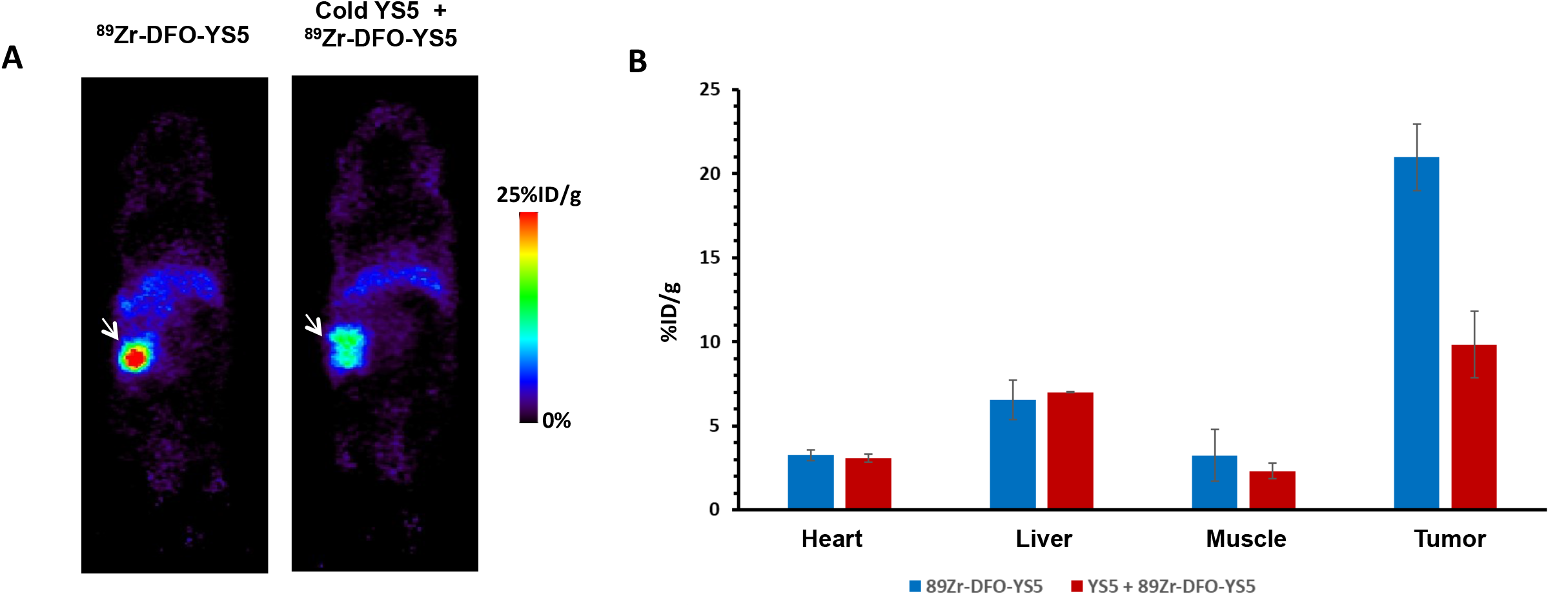
^89^Zr-DFO-YS5 PET imaging of a prostate cancer PDX model. (A) PET images of PDX, with (right) or without (left) competition with cold YS5 antibody. (B) SUV values derived from PET images for tumor and normal organs (hear, liver and muscle). Blue bar: SUV derived from ^89^Zr-DFO-YS5 PET images without cold YS5. Orange bar: SUV derived from ^89^Zr-DFO-YS5 PET images with cold YS5.

**Fig. 8.**
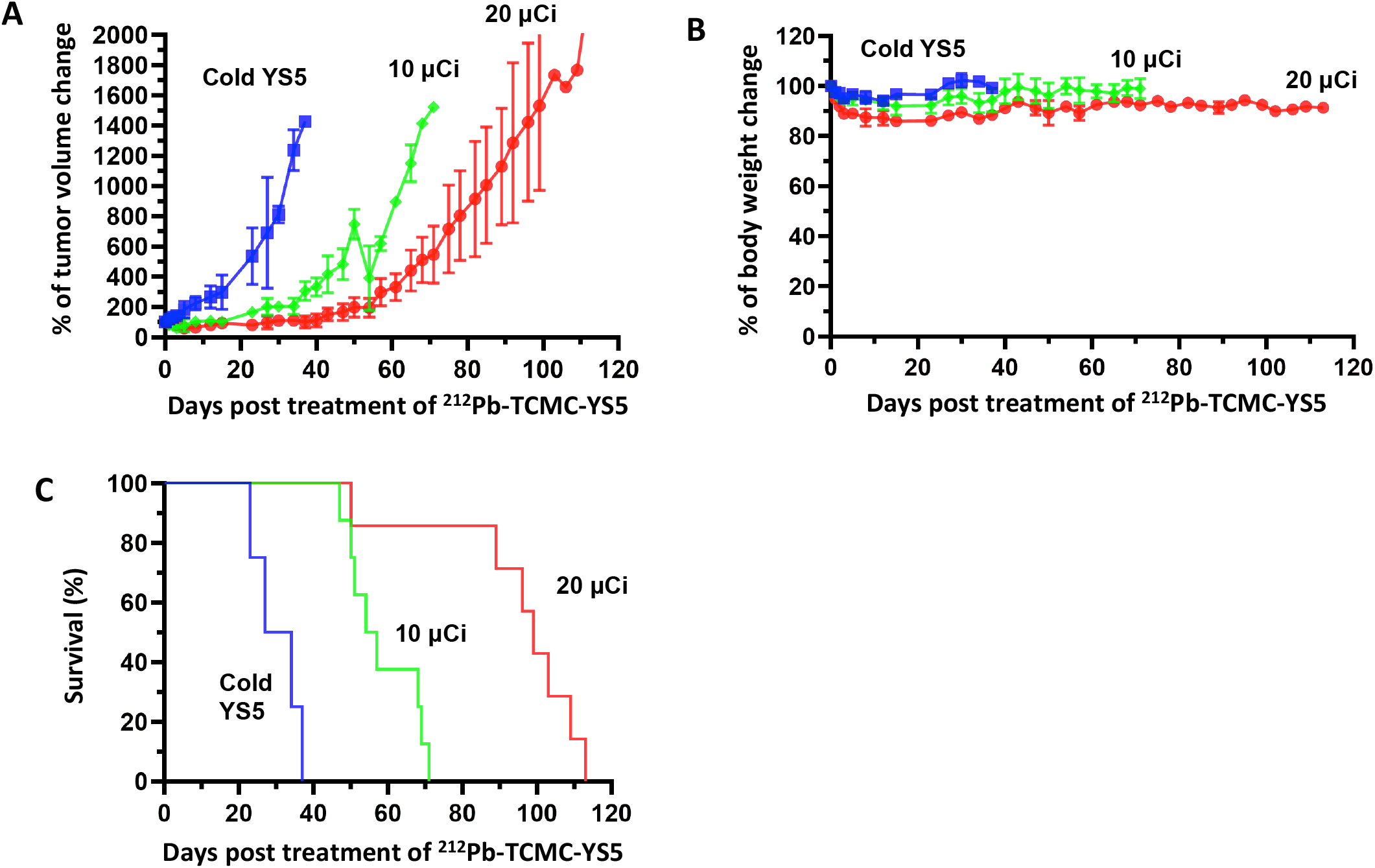
Therapeutic effect of a single dose of ^212^Pb-TCMC-YS5 in a prostate cancer PDX model. Blue: cold YS5 control. Green: a single dose of 10 μCi. Red: a single dose of 20 μCi. (A) Normalized tumor volume change (against day 0) post treatment. (B) Normalized body weight change (against day 0) post treatment. (C) Normalized animal survival rate (against day 0) post treatment.

## Discussion

CD46 is a new prostate cancer cell surface target that we identified and validated, which shows lineage independent homogeneous expression in both adenocarcinoma and small cell neuroendocrine mCRPC (15), differentiating from commonly targeted antigens such as PSMA. YS5, the human antibody that we developed and used to construct the CD46 targeting ADC (FOR46) that is in multiple clinical trials, binds to a tumor selective conformational epitope, enabling therapeutic targeting of CD46 (15,26). In this study we developed an YS5-based CD46-targeted ^212^Pb radiotherapy and studied efficacy in three prostate cancer animal models. We optimized the ^212^Pb labeling process that produced ^212^Pb-YS5 with high yield that retained high affinity binding to tumor cells. For therapeutic evaluation, we found that a single i.v. dose of 20 μCi of ^212^Pb-TCMC-YS5 was well tolerated without causing either acute or long-term toxicity as assessed by body weight change, general health condition, blood cell count and blood chemistry analysis. We next studied efficacy of this dose (a single injection) in three mouse models (i.e., subcutaneous and intraprostate orthotopic CDX models and a PDX model), and found potent tumor inhibition with an increase in animal survival in all three models.

Despite the relatively short half-life of ^212^Pb (10.6 h), the tumor inhibition effect from a single dose treatment lasted over 40 days post treatment, suggesting that ^212^Pb-TCMC-YS5 could be a promising candidate for prostate cancer therapy development. Several other studies also showed that ^212^Pb labeled full-length monoclonal antibodies were effective in tumor inhibition in animal models (4,5,27–29) and in early phase clinical trials (2,3), again supporting therapy development with ^212^Pb. Thus, ^212^Pb radio-ligand therapy may be a promising approach not only for small molecule ligands (8) but also large molecules such as a full-length monoclonal antibody including an IgG molecule. With the recently improved production of ^224^Ra-^212^Pb and ^203^Pb as the imaging theranostic pair, the pace of ^212^Pb-based radio-ligand therapy development is expected to accelerate (30–33).

It should be noted that the ^212^Pb radiolabeling process employed in this study can be readily incorporated into an automatic synthesizer for clinical production. ^212^Pb eluted by strong acids from the ^224^Ra-^212^Pb generator requires further process before labeling (20,29). The short half-life (10.6 h) of ^212^Pb also puts a time constraint on the preparation protocol. As such, a more facile and rapid process may facilitate clinical translation. Pre-concentration of ^212^Pb via extraction chromatography resin was reported recently (21), bypassing the strong acid requirement. A similar approach for peptide radiolabeling has been developed into a cassette-based system customized for clinical radiopharmaceutical production by the Schultz group (21). We adopted and modified the above methods, enabling completion of the whole labeling process for ^212^Pb-TCMC-YS5 within 2 h, suggesting that this method of ^212^Pb-TCMC-YS5 preparation could be further scaled for clinical production.

We studied the therapeutic effect of ^212^Pb-TCMC-YS5 in two mCRPC CDXs and one PDX – in all three models efficacy is evident for a single dose (20 μCi) and a single treatment cycle, with excellent tolerability and no clear toxicity detected by blood analysis 90 days post treatment. A single treatment of a lower dose (10 μCi) of ^212^Pb-TCMC-YS5 also showed efficacy in the PDX model, suggesting opportunities for further optimization. Additional dosing schemes and multiple treatment cycles in future studies could further improve efficacy over toxicity and increase the therapeutic window. Finally, the clinical stage human antibody YS5 is compatible with the development of radioimmunotherapy based on other alpha particles. For example, we have also conjugated YS5 with ^225^Ac and found potent anti-tumor activity in other prostate cancer models (BioRxiv 2022.10.13.512165). Further studies will compare efficacy and toxicity of those YS5 alpha particle therapies to select a candidate for clinical testing in human patients.

## Conclusion

The result from this first of its kind CD46-targeted ^212^Pb radioimmunotherapy study demonstrates that ^212^Pb-TCMC-YS5 is well tolerated and efficacious against prostate cancer in both CDX and PDX models. Given that CD46 is becoming a credentialed target in mCRPC, an YS5-based antibody-drug conjugate is in multiple clinical trials (NCT03575819 and NCT05011188) with an established safety profile, and an YS5-PET imaging agent is in a first-in-human study (NCT05245006), our YS5-conjugated radioimmunotherapy has potential for rapid translation into the clinic for prostate cancer treatment.

## Acknowledgements

This study was supported by NIH/NCI R01CA223767 (JH and BL) and CCSG P30 CA44579 (University of Virginia Comprehensive Cancer Center). The PET imaging data and the bioluminescence data were acquired through the University of Virginia Molecular Imaging Core Lab, with NIH S10OD021672 funding for the Albira Si trimodal scanner. The CDX orthotopic model was performed through University of Virginia Molecular Assessments and Preclinical Studies (MAPS) Core. The isotope ^212^Pb used in this research was supplied by US Department of Energy Isotope Program managed by the Office of Isotope R&D and Production. The authors are grateful to Dr. Martin Brechbiel for providing advice on translational development, Drs. Michael Schultz and Mengshi Li for providing advice on the Pb resin, and Dr. Kevin Lynch at University of Virginia for the use of Heska Element HT5 hematology analyzer.

## Author contribution

J.L. performed *in vitro* cell studies and in vivo imaging and therapy studies on CDX subcutanouse model, analyzed the data.

T.H. performed *in vivo* studies on CDX orthotopic model and PDX model, analyzed the data.

J. Hua. performed *in vitro* cell studies and *in vivo* therapy studies on CDX subcutaneous model, analyzed the data.

Q.W. performed the PC3-luc tumor cell study.

Y.Su optimized the YS5 antibody purification process and participated in data analysis.

P.C. performed the HPLC analysis of radiolabeled antibodies

S.B. performed antibody purification and characterization, and data analysis.

A.L. performed antibody purification and characterization.

Z.X. performed cell culture of PC3-luc cells

A.B. provided *in vivo* therapy antibody dose guidance.

S.S. provided guidance on *in vivo* therapy and revised the manuscript.

W.S. provided guidance on *in vivo* therapy.

Y.S. provided experimental guidance.

R.R.F. provided experimental guidance.

D.G. provided experimental guidance and revise the manuscript.

R.D. provided experimental guidance and revise the manuscript.

H.L. provided experimental guidance and support on PC3-luc cell studies.

B.L. conceptualized the study, designed, funded, guided the project, and wrote the manuscript.

J.H. conceptualized the study, designed, funded, guided the project, and wrote the manuscript.

