## Supplemental tables and figures for "CD46 targeted ^212^Pb alpha particle radioimmunotherapy for prostate cancer treatment"

**List of supplemental data:**

**Table S1:** Hematology parameters of acute (day-14) and long-term (day-90) toxicity studies.

**Table S2:** Blood clinical chemistry tests for liver and kidney function for acute and long-term toxicity studies.

**Figure S1:** Therapeutic effects of a single dose $^{212}$Pb-TCMC-YS5 in the PC3 subcutaneous CDX model.

**Figure S2:** BLI images of the PC3-Luc orthotropic CDX model treated by a single dose $^{212}$Pb-TCMC-YS5.

**Figure S3:** Change of actual tumor volume of individual PDX mouse treated with a single dose of 10 and 20 μCi of $^{212}$Pb-TCMC-YS5.

**Table S1**

Hematology parameters of acute (day-14) and long-term (day-90) toxicity studies. RBC: Red blood cell; WBC: White blood cell; PLT: Platelet; HCT: Hematocrit test (proportion of RBC in blood); HGB: Hemoglobin; Neu: Neutrophil; Lym: Lymphocyte; Mon: Monocyte; and Eos: Eosinophil. Mean ± SD, n = 5

| | 14 days | | 90 days | |
|---|---|---|---|---|
| | $^{212}$Pb-TCMC-YS5 | No treatment | $^{212}$Pb-TCMC-YS5 | No treatment |
| RBC ($10^6$/µL) | 7.62±0.66 | 7.99±0.23 | 6.68±0.57 | 8.15±0.08 |
| WBC ($10^3$/µL) | 4.15±0.94 | 5.28±1.72 | 6.23±3.07 | 5.39±1.29 |
| PLT ($10^3$/µL) | 1366.50±208.76 | 1515.20±107.71 | 1494.83±219.88 | 1625.00±160.97 |
| HCT | 35.98±3.51 | 39.52±1.14 | 31.47±4.13 | 39.02±1.46 |
| HGB (g/dL) | 12.58±1.24 | 13.48±0.49 | 11.05±1.11 | 13.18±0.56 |
| Neu (%) | 26.35±9.38 | 13.40±1.94 | 17.98±3.80 | 19.44±5.74 |
| Lym (%) | 68.35±11.45 | 82.62±3.36 | 76.13±6.11 | 74.80±8.10 |
| Mon (%) | 4.02±1.87 | 1.62±0.90 | 2.92±1.60 | 2.44±1.79 |
| Eos (%) | 1.23±0.42 | 2.34±1.05 | 2.88±1.86 | 3.26±1.46 |

**Table S2**

Blood clinical chemistry tests for liver and kidney function for acute and long-term toxicity studies. Mean ± SD, n = 5.

| | 14 days | | 90 days | |
|---|---|---|---|---|
| | $^{212}$Pb-TCMC-YS5 | No treatment | $^{212}$Pb-TCMC-YS5 | No treatment |
| Albumin (µg/mL) | 34.8±3.8 | 26.4±10.9 | 30.1±5.5 | 30.4±5.3 |
| Creatinine (µmol/L) | 18.7±5.6 | 15.1±5.8 | 17.3±1.2 | 14.7±1.4 |
| Blood Urea Nitrogen (mmol/L) | 15.8±3.2 | 9.4±4.1 | 13.3±4.6 | 13.2±4.6 |
| Alanine transaminase (ng/mL) | 15.9±2.0 | 13.6±0.9 | 16.3±1.1 | 15.5±1.2 |
| Aspartate transaminase (ng/mL) | 113.5±11.3 | 91.4±32.6 | 119.3±29.3 | 106.3±29.7 |

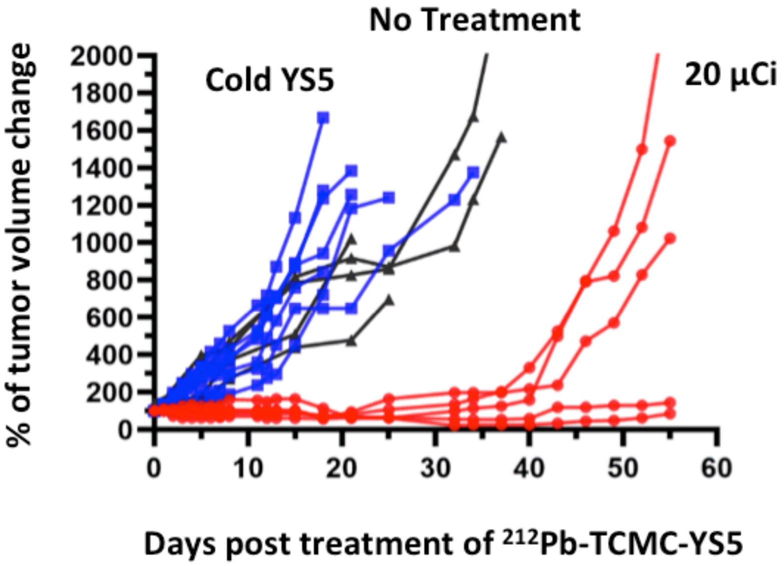


**Figure S1:** Therapeutic effects of a single dose $^{212}$Pb-TCMC-YS5 in the PC3 subcutaneous CDX model. Tumor growth of individual mice within 55 days post treatment of $^{212}$Pb-TCMC-YS5 (normalized against tumor volume at day 0). Red: Study group receiving $^{212}$Pb-TCMC-YS5. Blue: Control group receiving unlabeled YS5 antibody only.

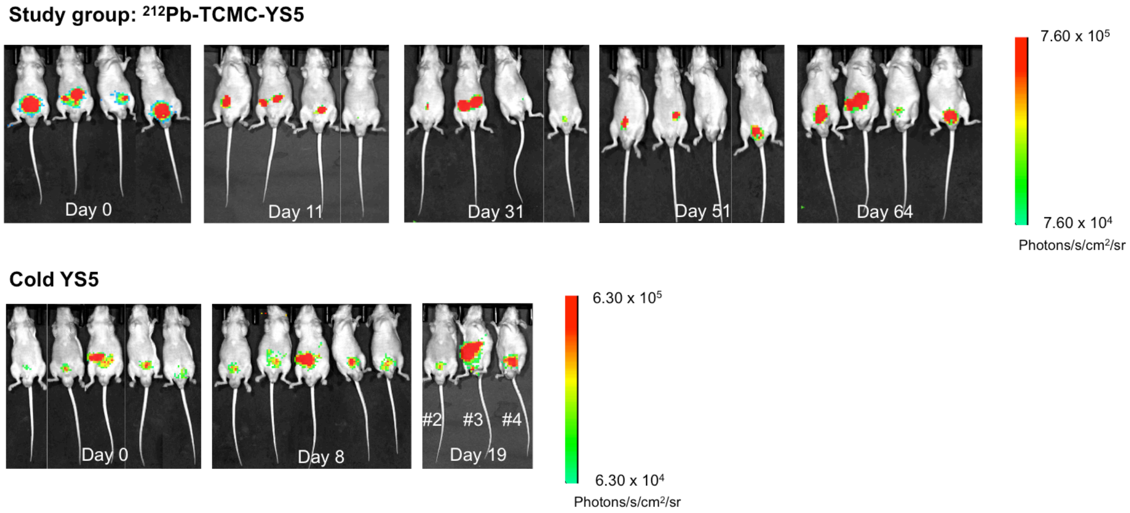


**Figure S2:** BLI images of the PC3-Luc orthotropic CDX model treated by a single dose $^{212}$Pb-TCMC-YS5 (top), and the cold YS5 control (bottom). Two animals died in the control group on day-19 post treatment and were not imaged.

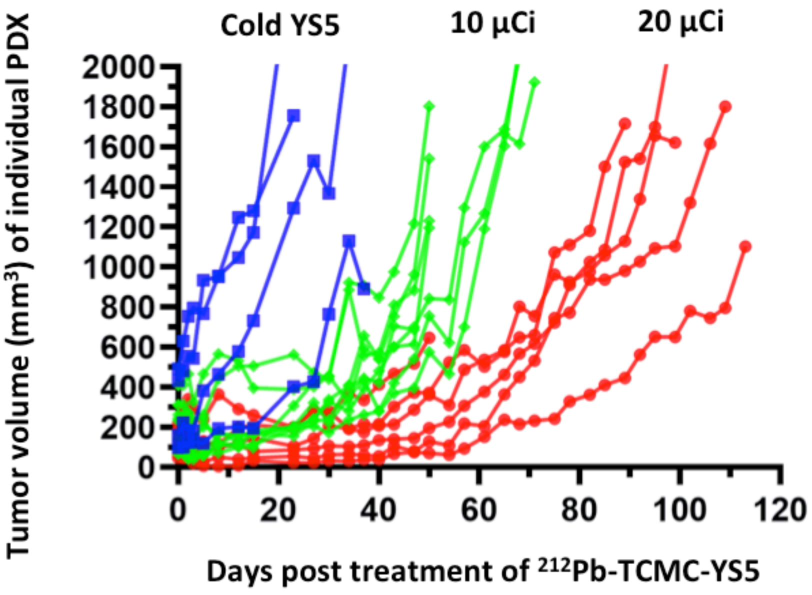


**Figure S3:** Change (days post injection) of actual tumor volume of individual PDX mouse treated with a single dose of 10 and 20 µCi of $^{212}$Pb-TCMC-YS5.
